# Metal Ion-Releasing Glass Particles to Enhance Antibiotic Efficacy Against Cystic Fibrosis Infection

**DOI:** 10.1101/2025.10.30.685604

**Authors:** Maxwell M. Wolverton, Zhenyu Cheng, Daniel Boyd, Brendan M. Leung

## Abstract

Cystic fibrosis is characterized by thickened airway mucus that impairs mucociliary clearance and promotes persistent bacterial infections. These chronic infections, primarily caused by *Pseudomonas aeruginosa* and *Staphylococcus aureus*, lead to progressive lung damage and respiratory failure, which are the leading causes of mortality in cystic fibrosis patients. Although antibiotics remain the cornerstone of treatment for such airway infections, incomplete bacterial eradication often contributes to the emergence of antibiotic resistance. This study explores whether the efficacy of conventional antibiotics can be enhanced through co-delivery with antibacterial metal ions released from borate-based bioactive glasses. To evaluate this combination therapy, we developed an *in vitro* airway infection model that replicates key features of the cystic fibrosis airway environment. The model incorporates a layer of bronchial epithelial cells, a mucus-like hydrogel, and bacteria deposited using an aqueous two-phase system. Multiple bioactive glass formulations were evaluated for their ability to augment antibiotic activity and enhance bacterial eradication. The results demonstrated additive or synergistic antibacterial effects against *P. aeruginosa* and *S. aureus*, while maintaining mammalian cell viability. These findings suggest that metal ion-antibiotic combination therapies delivered via bioactive glasses may improve treatment outcomes for cystic fibrosis-related infections and reduce reliance on high-dose or prolonged antibiotic regimens. Such approaches hold promise not only for cystic fibrosis patients but also for broader clinical applications where antibiotic resistance is a growing concern.

## Introduction

Lung damage and respiratory failure caused by chronic airway infections are the leading causes of mortality for patients with cystic fibrosis (CF) [1], [2]. Infections of the CF airway occur due to the compromised microenvironment created by the altered airway mucus layer of CF [3]. In CF pathology, mutations in the cystic fibrosis transmembrane conductance regulator (*CFTR*) gene lead to a reduction in water transport across the airway epithelium, resulting a dehydrated and more viscous airway mucus layer [3], [4], [5]. These properties of CF mucus lead to increased airway colonization by opportunistic pathogens because of impairment to the mucociliary clearance mechanism, host immune cell migration, and clearance by coughing [3], [5], [6].

*Pseudomonas aeruginosa* and *Staphylococcus aureus* are the predominant bacteria responsible for airway infections in CF patients [7], [8]. Antibiotic therapy is the primary form of treatment for such CF airway infections, aiming to reduce microbial load and slow lung damage; however, the widespread use of antibiotics has contributed to rising antibiotic resistance in CF pathogens [1], [9], [10]. In a CF context, antibiotic resistance develops as bacteria survive sublethal antibiotic exposure within the thick, mucus-filled airways, where impaired drug diffusion and biofilm formation create an environment that promotes the persistence and spread of resistant strains [10], [11]. Infections caused by drug-resistant pathogens are currently responsible for an estimated 1.2 million deaths each year, with projections suggesting this number could rise to 10 million annually by 2050 if effective measures are not implemented to control antibiotic resistance [12], [13]. This highlights the critical need for innovative therapeutic strategies that not only treat infections effectively but also minimize the emergence and spread of antibiotic resistance.

This study aims to enhance the treatment of CF bacterial airway infections by combining conventional antibiotics with antibacterial metal ions to target bacteria through multiple complementary mechanisms. Metal ions exhibit antimicrobial activity through multiple mechanisms, including membrane and DNA damage, enzyme inhibition, and the generation of reactive oxygen species (ROS), with ROS playing a key role in both antibacterial and antibiofilm effects [14], [15]. Metal ions target microbes through multiple simultaneous mechanisms, reducing the likelihood of antibiotic resistance development [16], [17]. Therefore, combining metal ions with conventional antibiotics will expand the range of antibacterial actions, further minimizing the potential for resistance to emerge [17]. In the context of CF airway infections, metal ions offer added benefits due to their ability to penetrate thick mucus and potentially disrupt mucus structure through mechanisms such as ROS generation [17], [18]. This study examined zinc and gallium ions, with zinc shown to disrupt metabolism and generate ROS, and gallium disrupting iron metabolism to kill pathogens such as *P. aeruginosa* and *M. tuberculosis* [16], [19].

While metal ions have potential for treating CF airway infections, an effective delivery system is necessary to transport them to sites of infection. Borate-based bioactive glasses have shown promise as degradable carriers for metal ions and controlled drug delivery in various medical applications, including bone regeneration, chronic infection treatment, wound healing, and cancer therapy [20], [21], [22]. Their highly tunable degradability makes them well-suited for delivering antibacterial metal ions to airway infection sites while ensuring complete dissolution [23].

A representative CF infection model is essential for evaluating the antibacterial effectiveness of new therapies under conditions that mimic the CF airway environment. For this study, we designed an *in vitro* airway infection model with mammalian cells, mucus-mimetic hydrogels, and common CF bacteria deposited using an aqueous two-phase system (ATPS). This infection model builds upon the model designed by O’Brien et al. [24]. A key component of the model is the mucus-mimetic hydrogels composed of an interpenetrating alginate-mucin (ALG-MUC) network crosslinked with CaCl_2_. Two formulations were created to represent healthy and CF-like mucus, with the CF formulation containing higher levels of crosslinker and mucin to reflect the thicker, more viscous properties of CF airway secretions [24], [25]. The newly designed model utilizes a transwell insert system to more accurately mimic the *in vivo* airway environment and allow for easier assayability.

The primary objective of this study was to evaluate if incorporating metal ion-releasing bioactive glass could enhance the effectiveness of conventional antibiotics in eradicating bacterial infections within a CF-like airway microenvironment. To accomplish this objective, the airway infection model was cultured with combinations of conventional antibiotics with metal ion-releasing bioactive glass extracts, followed by assessment of antibacterial efficacy and mammalian cell cytotoxicity (Figure 1). The findings of this study demonstrate the potential of combining metal ions with conventional antibiotics as an effective treatment for CF airway infections, while also helping to reduce the risk of resistance development.

**Figure 1.**
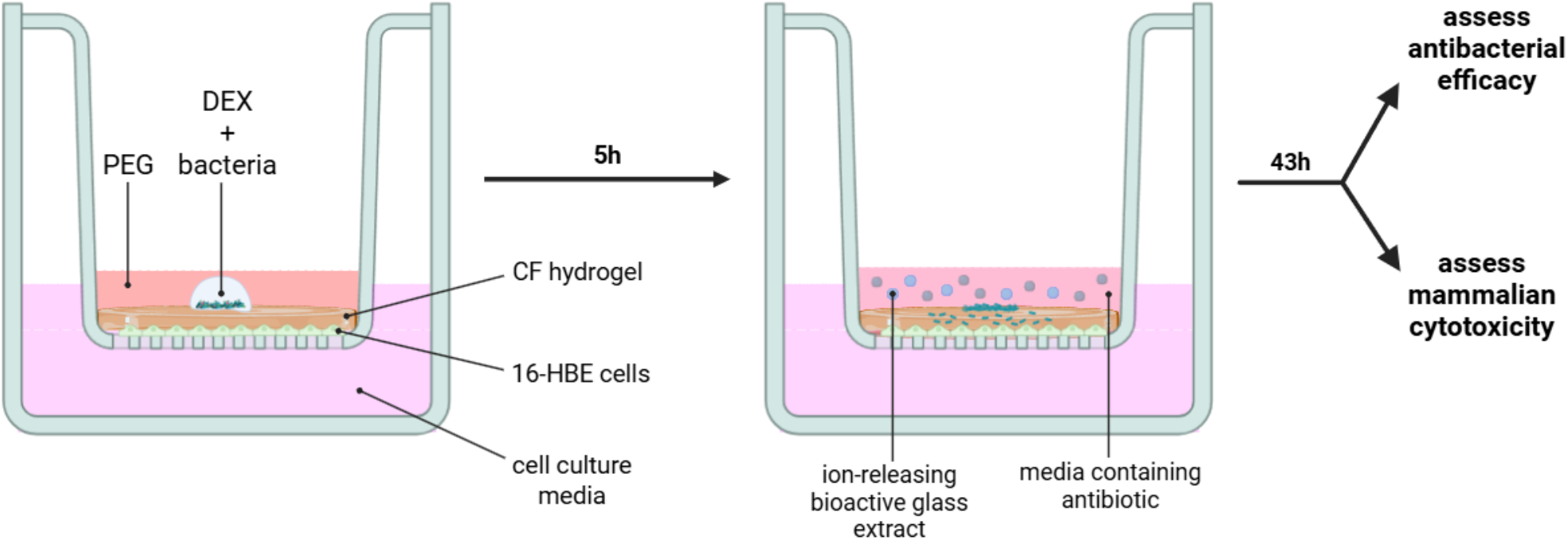
Evaluation of efficacy and safety of metal ion-releasing bioactive glasses combined with conventional antibiotics. The *in vitro* airway infection model consists of bronchial epithelial cells (16-HBE) overlaid with a CF-mimetic hydrogel layer and bacteria using an aqueous two-phase system (ATPS) composed of polyethylene glycol (PEG) and dextran (DEX). The liquid phase of the model is replaced with media containing the antibiotic-glass extract combinations five hours after bacteria deposition. The model is then cultured for an additional 43 hours (total 48 hours) before evaluating the antibacterial efficacy of the antibiotic-glass combinations and their impact on mammalian cell viability. Diagram created with BioRender.com.

## Methods

### Mammalian cell culture conditions

The 16-HBE cell line (16HBE14o-), kindly provided by Dr. Geoff Maksym from Dalhousie University, was used in the airway infection model to mimic the mammalian bronchial epithelial cell layer. 16-HBE cells were incubated at 37 °C and 5% CO_2_ and were cultured for growth using a 1:1 ratio of Dulbecco’s Modified Eagle Medium (DMEM), high glucose (Thermo Fisher Scientific, Gibco, Cat. No. 11965092), and Gibco Ham’s F-12 Nutrient Mixture (Thermo Fisher Scientific, Gibco, Cat. No. 11765054) supplemented with 10% fetal bovine serum (FBS; Thermo Fisher Scientific, Gibco, Cat. No. A5456701) and 1% antibiotic-antimycotic (AA; Thermo Fisher Scientific, Gibco, Cat. No. 15240062). In the airway infection model, the 16-HBEs were cultured with a 1:1 ratio of DMEM, low glucose (Thermo Fisher Scientific, Gibco, Cat. No. 11885084), and F-12 supplemented with 1% FBS in order to slow 16-HBE proliferation and promote a quiescent-like state in the model.

### Bacteria culture conditions

*P. aeruginosa* CF18 and *S. aureus* ATCC 6538 were used in the airway infection model. *P. aeruginosa* CF18 was kindly provided by Dr. Zhenyu Cheng from Dalhousie University and was originally isolated from a child (<24 months) with CF [26]. Luria-Bertani (LB; Sigma-Aldrich, Cat. No. 1.10285) broth was used to culture *P. aeruginosa* while Brain Heart Infusion (BHI; Sigma-Aldrich, Cat. No. 1.10493) broth was used to culture *S. aureus*. The two bacteria strains were stored in their respective broths with 25% (v/v) glycerol at -80 °C. In preparation for an experiment, the frozen stocks were streaked on 1.5% (w/v) agar (Sigma-Aldrich, Cat. No. A1296) containing their respective broth. After incubating the streaked plates for 16 to 18 h at 37 °C and 5% CO_2_, liquid cultures were prepared by inoculating 3 ml of the respective broth with a single colony from the streaked plates. The inoculated broth was then incubated for 16 to 18 h at 37°C and 5% CO_2_ on a shaking incubator at 200 rpm prior to use for experiments.

### Aqueous two-phase system preparation

ATPS formulations in this study incorporated polyethylene glycol (PEG) and dextran (DEX) as phase-forming polymers. ATPS solutions were prepared by dissolving 5% (w/v) PEG (35 kDa,

Sigma-Aldrich, Cat. No. 8.18892) and 5% (w/v) DEX (500 kDa, Pharmacosmos, Cat. No. 5510 0070 9006) in either DMEM/F-12 supplemented with 10% FBS or in LB broth, using a rocking platform shaker. The solutions were then sterile filtered and centrifuged at 3000 g for 90 min to sperate the two phases. The PEG-rich and DEX-rich phase of each ATPS solution was then collected and stored at 4 °C. In the airway infection model, the PEG-rich phase of the DMEM/F-12 solution was used to support mammalian cell growth, while the DEX-rich phase of the LB broth solution was used to support bacterial growth.

### Preparation of hydrogel stock solutions

Mucus-mimetic hydrogels were composed of an alginate-mucin interpenetrating network crosslinked by calcium chloride (CaCl_2_). Alginic acid sodium salt power (low viscosity, MP Biomedicals, Cat. No. 154725) was UV sterilized and then mixed with HEPES buffer to produce a 75 mM alginate stock solution. The CaCl_2_ stock solution was prepared by dissolving 2 mg/ml CaCl_2_ (Sigma-Aldrich, Cat. No. 746495) in deionized water and filter sterilized. To prepare the mucin stock solution, Mucin Type II powder (from porcine stomach, Sigma-Aldrich, Cat. No. M2378) was sterilized by saturation with 95% ethanol and then dissolved in PBS to produce a 200 mg/ml stock solution. Two hydrogel formulations were produced in this research to mimic healthy and CF mucus properties: a healthy hydrogel containing 1.5% (w/v) alginate, 0.5 mg/ml CaCl_2_, and 1% mucin, and a CF hydrogel containing 1.5% (w/v) alginate, 0.6 mg/ml CaCl_2_, and 4% mucin. The CF hydrogel formulation was used in the airway infection model to assess antibiotic-glass efficacy in a CF context.

### Airway infection model assembly

16-HBE cells were first seeded onto transwell inserts (VWR, Avantor, Cat. No. 734-3263) in a 24-well plate at a cell density of approximately 30000 cells/insert. To encourage cell adhesion, the inserts were coated with 0.05 mg/ml collagen (Advanced BioMatrix, BICO, PureCol® Bovine Collagen Solution, Type I, Cat. No. 5005) prior to cell seeding. Before assembling the model, the 16-HBE cells were grown in DMEM, high glucose/F-12 + 10% FBS for 48 hours to reach confluency, and then an additional 48 hours in DMEM, low glucose/F-12 + 1% FBS to induce a quiescent-like state.

After this time, the mucus-mimetic hydrogels were added to the model. First, a thin layer of an alginate-CaCl_2_ hydrogel formulation containing 0% mucin was deposited directly on top of the mammalian cells at a thickness of 0.4 mm. This alginate-only hydrogel was incorporated to improve cell adhesion by mitigating direct contact between the mammalian cell layer and the overlying mucin hydrogel. Then, the CF hydrogel formulation was deposited on top of the alginate-only hydrogel to produce a total hydrogel thickness of 1.5 mm to mimic physiological mucus layer thickness. To deposit the hydrogels into the model, the alginate and mucin solutions were first mixed together, then mixed with the CaCl_2_ solution and pipetted into the model. Immediately after deposition, the plate was centrifuged at 100 g for 5 minutes to improve hydrogel uniformity on the insert surface. The plate was then incubated at 37 °C and 5% CO_2_ to solidify: 10 minutes for the thin alginate-only hydrogel, and 30 minutes for the CF hydrogel.

After the hydrogels solidified in the incubator, the PEG-rich phase from a 5% PEG/5% DEX ATPS in DMEM/F-12 + 10% FBS was added to the apical compartment, covering the hydrogels. Using the BioMek 4000 Automated Liquid Handler (Beckman Coulter), a 0.5 µl droplet of the DEX-rich phase from a 5% PEG/5% DEX ATPS in LB broth containing the selected bacteria at an OD_600_ of 0.01 was carefully dispensed directly above the hydrogel layer within the PEG-rich phase. The model was then incubated at 37 °C and 5% CO_2_ for 5 hours to allow for controlled bacterial settling into the hydrogel phase. After 5 hours, the liquid phase was then removed and replaced with media containing the antibiotic-glass combinations, and then the model was incubated at 37 °C and 5% CO_2_ for an additional 43 hours, bringing the total culture time to 48 hours. After incubation, bacterial growth and cell viability were evaluated.

### Preparation of antibiotic-glass combinations

The ionic compositions of the three bioactive glass formulations selected for this study are shown in Table 1. The three glasses selected for this study were chosen based off of antibacterial performance against planktonic *P. aeruginosa* and *S. aureus* with minimal mammalian cell cytotoxicity. For each glass extract, 0.1 g of each glass was added to 10 ml of cell culture media in a 50 ml Falcon tube and incubated at 37 °C with shaking at 200 rpm for 48 hours to promote glass dissolution and metal ion release. The resulting mixture was centrifuged, and the supernatant—containing the dissolved ions—was collected and labelled as “dilution 1.” This stock extract was then further diluted to the appropriate concentrations for each experiment.

**Table 1.** Compositions of the three bioactive glass formulations used in this study.

| Glass Identifier | B<sub>2</sub>O<sub>3</sub><br>mol % | Na<sub>2</sub>O<br>mol % | ZnO<br>mol % | Ga<sub>2</sub>O<sub>3</sub><br>mol % |
| --- | --- | --- | --- | --- |
| Ga<sup>3+</sup> Glass | 56.28 | 30 | 0 | 13.72 |
| Zn<sup>2+</sup> Glass | 62.78 | 30 | 7.22 | 0 |
| Ga<sup>3+</sup>/Zn<sup>2+</sup> Glass | 50 | 25.90 | 4.10 | 20 |

### Assessment of bacterial growth in the airway infection model

For single-strain experiments, bacterial growth in both the liquid and hydrogel phases was quantified using spot plating to determine CFU/ml. A 20 µl sample from the liquid phase was collected, while the remaining liquid was discarded. The hydrogel phases were transferred to microcentrifuge tubes, centrifuged at 10,000 g for 10 minutes to pellet bacteria, and resuspended in 50 µl of PBS. Both liquid and hydrogel samples were serially diluted (10^1^-10^6^) in a 96-well plate, and 20 µl from each dilution was spot plated onto sectioned LB agar plates. After incubating at 37 °C and 5% CO_2_ for 16-18 hours, colonies were counted from dilutions with 30-300 colonies. CFU/ml was then calculated using eq 1, where N is the viable bacteria concentration, C is the colony count, V is the plated volume in ml, and D is the dilution factor.

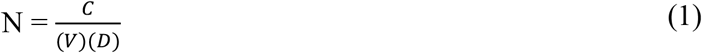

For co-culture experiments with *P. aeruginosa* and *S. aureus*, spot plating was not suitable due to indistinguishable colony morphology in small droplets. Instead, spread plating was used to differentiate and quantify each species. Samples were diluted as before, and 100 µl of a single dilution was spread across an LB agar plate using a sterile spreader. This allowed for a clear distinction between *P. aeruginosa* (blue, fuzzy colonies) and *S. aureus* (yellow, sharp colonies) based on morphology and color. CFU/ml for each species was then determined using eq 1.

### Assessment of mammalian cell viability in the airway infection model

A live/dead assay consisting of calcein AM (Thermo Fisher Scientific, Gibco, Cat. No. C1430) and ethidium homodimer-1 (Thermo Fisher Scientific, Gibco, Cat. No. E1169) and a Hoechst stain (Hoechst 33342, Thermo Fisher Scientific, Gibco, Cat. No. H3570) was performed to assess the mammalian cell viability in the airway infection model after being cultured with the antibiotic-glass combinations. The live/dead assay was performed without disturbing the apical compartment to preserve model structure and retain unattached dead cells. Staining solutions were sequentially added to the basolateral compartment, followed by incubation and rinsing steps, before imaging in PBS. The EVOS™ FL Auto 2 Imaging System (Thermo Fisher Scientific) was then used to capture images of live cells (green fluorescent protein channel, GFP), dead cells (red fluorescent protein channel, RFP), and DNA (DAPI channel). ImageJ was used to quantify dead-stained and Hoechst-stained cells, and these counts were used to calculate cell viability according to eq 2.

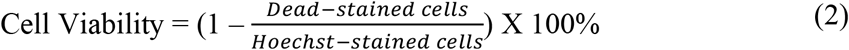

### Statistical analysis

GraphPad Prism (Version 10.3.1) was used for all statistical analyses. An ordinary two-way analysis of variance (ANOVA), followed by Šidák’s multiple comparisons test, was used to assess statistical significance of the antibacterial efficacy data for the antibiotic-glass combinations. p-values were reported as follows: *p < 0.05, **p < 0.01, ***p < 0.001, ****p < 0.0001. This analysis included two factors: glass dilution (three levels, specific to each experiment) and antibiotic concentration (two levels: 0 and 16 µg/ml). This analysis allowed us to determine the statistically significant effects of glass, both alone and combined with antibiotic, on bacterial growth in the airway infection model.

## Results

### *P. aeruginosa* and *S. aureus* growth is inhibited by certain antibiotic-glass combinations in the monomicrobial CF airway infection model

Each combination of antibiotic (tobramycin or vancomycin) with one bioactive glass extract (Ga^3+^, Zn^2+^, or Ga^3+^/Zn^2+^) was added to the CF airway infection model inoculated with either *P. aeruginosa* or *S. aureus* for a total of 48 hours. Following treatment, bacterial growth was quantified using a CFU/ml assay.

*P. aeruginosa* growth in the CF airway infection model was significantly decreased in the presence of the Ga^3+^ and Ga^3+^/Zn^2+^ glasses compared to the no treatment control (Figure 2). *P. aeruginosa* growth was not significantly affected by the presence of the Zn^2+^ glass (Figure 2). The combination of Ga^3+^ glass with tobramycin resulted in reduced *P. aeruginosa* growth within the model compared to either treatment alone, suggesting that Ga^3+^ glass enhances the antibacterial efficacy of tobramycin (Figure 2). In contrast, combining tobramycin with the Ga^3+^/Zn^2+^ glass did not further reduce *P. aeruginosa* growth compared to the glass alone, suggesting no additional benefit from the combination (Figure 2).

**Figure 2.**
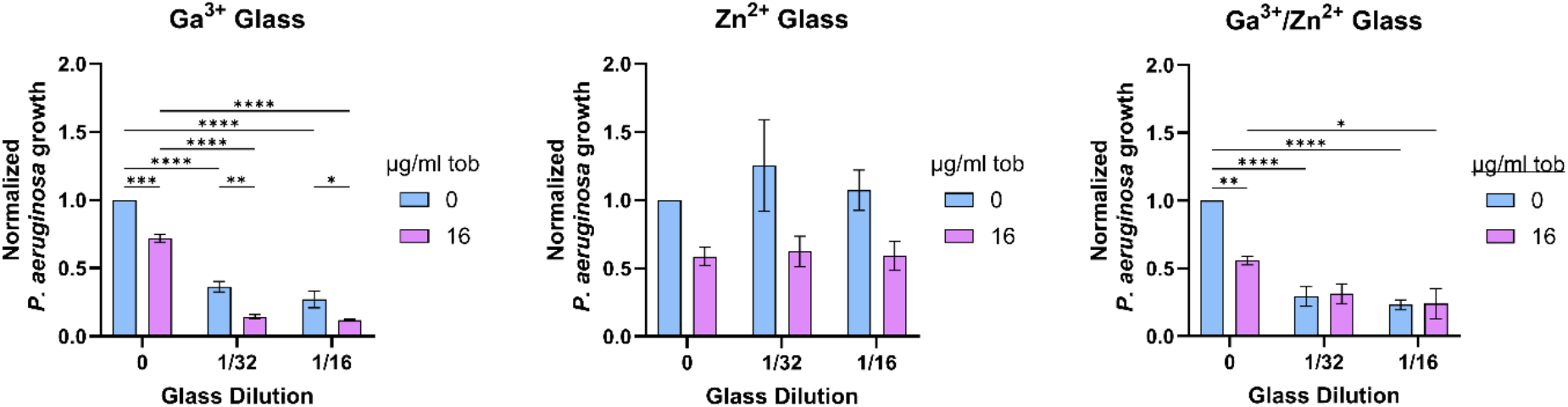
*P. aeruginosa* growth in the hydrogel phase of the CF airway infection model following treatment with various tobramycin-glass combinations. Models were treated for 48 hours with bioactive glass extracts containing Ga^3+^ and/or Zn^2+^ ions at two dilutions (1/32, 1/16), with or without 16 µg/ml tobramycin. Bacterial growth was normalized to the untreated control. Data are presented as mean ± SEM (N = 3). Statistical analysis was performed using two-way ANOVA with Šidák’s multiple comparisons test; * p < 0.05, ** p < 0.01, *** p < 0.001, **** p < 0.0001.

*S. aureus* growth in the CF airway infection model was significantly decreased in the presence of the Ga^3+^ and Zn^2+^ glasses compared to the no treatment control (Figure 3). However, *S. aureus* growth was not significantly reduced in the presence of the Ga^3+^/Zn^2+^ glass (Figure 3). The combination of the Zn^2+^ glass at the 1/4 dilution with vancomycin led to reduced *S. aureus* growth in the hydrogel phase compared to either treatment alone, indicating that the Zn^2+^ glass may enhance the antibacterial activity of vancomycin (Figure 3). In contrast, combining vancomycin with the Ga^3+^ glass did not further reduce *S. aureus* growth relative to the Ga^3+^ glass alone, suggesting no additional suppressive effect from the combination (Figure 3).

**Figure 3.**
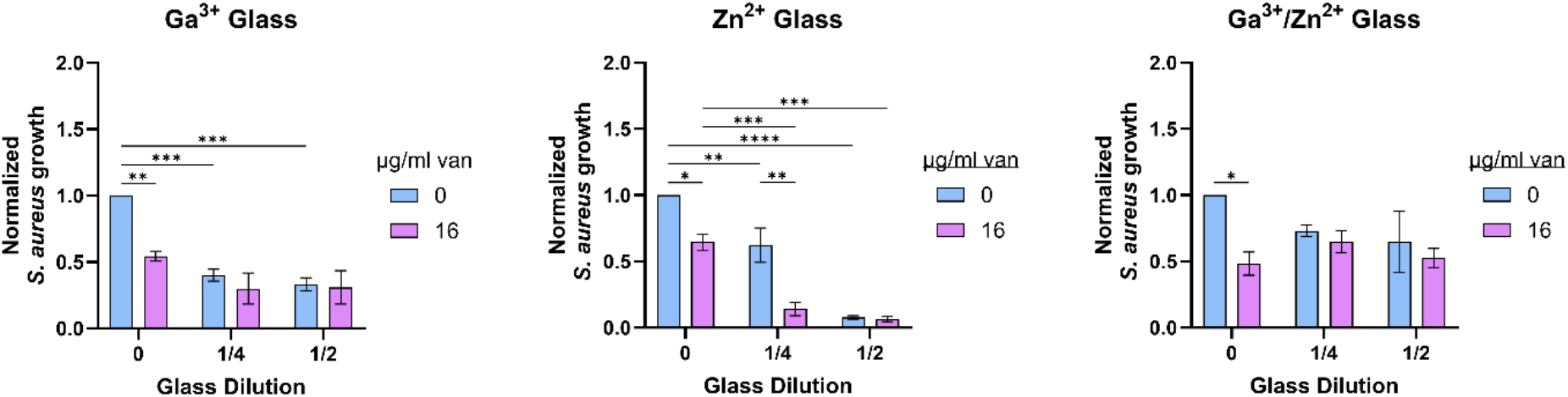
*S. aureus* growth in the hydrogel phase of the CF airway infection model following treatment with various vancomycin-glass combinations. Models were treated for 48 hours with bioactive glass extracts containing Ga^3+^ and/or Zn^2+^ ions at two dilutions (1/4, 1/2), with or without 16 µg/ml vancomycin. Bacterial growth was normalized to the untreated control. Data are presented as mean ± SEM (N = 3). Statistical analysis was performed using two-way ANOVA with Šidák’s multiple comparisons test; * p < 0.05, ** p < 0.01, *** p < 0.001, **** p < 0.0001.

### *P. aeruginosa* and *S. aureus* growth is inhibited by certain antibiotic-glass combinations in the polymicrobial CF airway infection model

Each combination of antibiotic (tobramycin or vancomycin) with one bioactive glass extract (Ga^3+^, Zn^2+^, or Ga^3+^/Zn^2+^) was added to the polymicrobial CF airway infection model inoculated with both *P. aeruginosa* and *S. aureus* for a total of 48 hours. Following treatment, bacterial growth was quantified using a CFU/ml assay to assess how bacterial growth responses to the antibiotic-glass combinations differ in a co-infection context.

When tobramycin-glass combinations were added to the polymicrobial CF airway infection model, treatment with the Ga^3+^ or the Ga^3+^/Zn^2+^ glass resulted in a significant reduction in *P. aeruginosa* growth within the hydrogel phase compared to the no treatment control (Figure 4). In contrast, the Zn^2+^ glass did not produce a significant change in *P. aeruginosa* growth (Figure 4). *S. aureus* growth in the polymicrobial model was increased in the presence of the Ga^3+^ and the Ga^3+^/Zn^2+^ glass compared to the no treatment control, while the Zn^2+^ glass did not produce a significant change in *S. aureus* growth (Figure 4).

**Figure 4.**
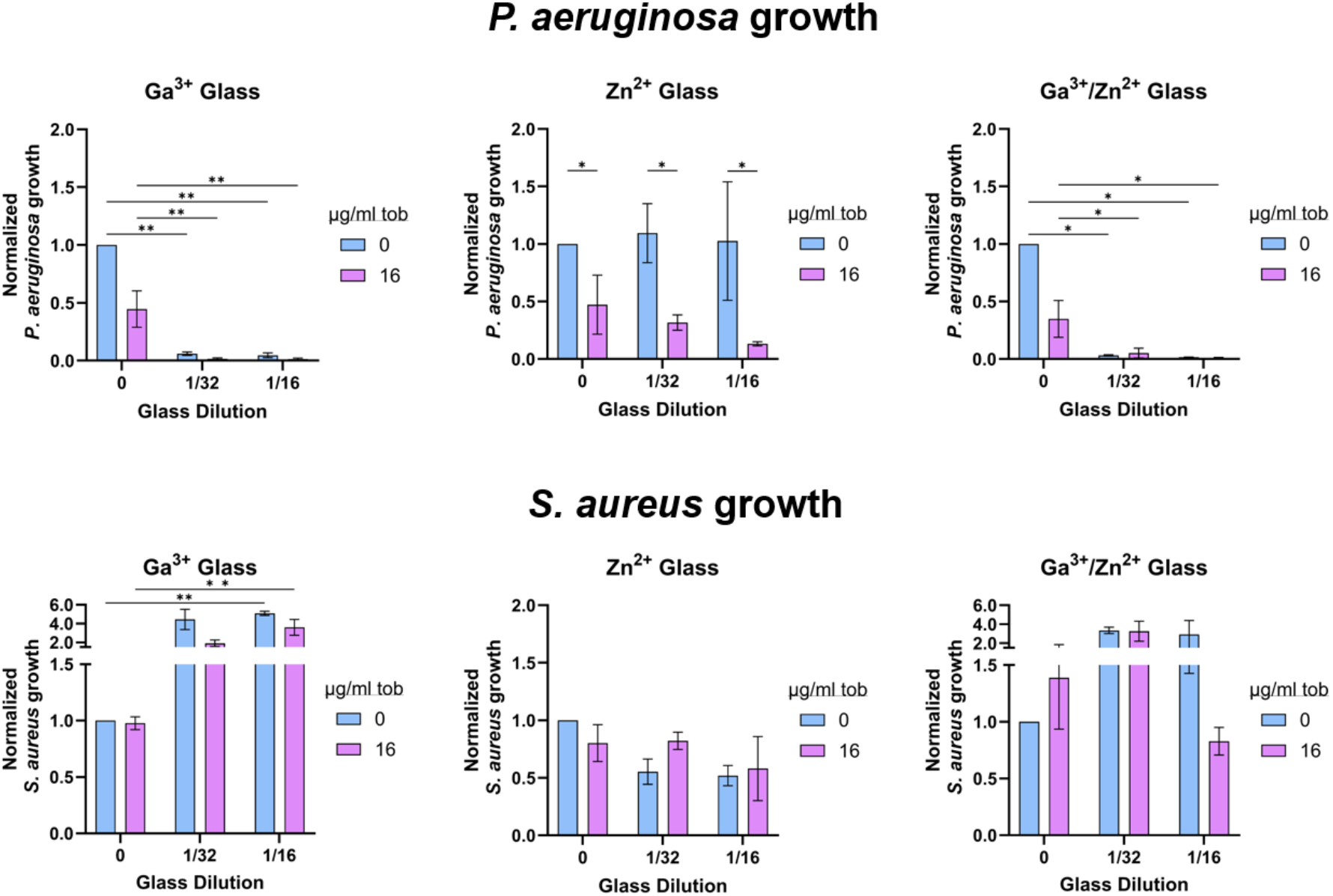
*P. aeruginosa* growth and *S. aureus* growth in the hydrogel phase of the polymicrobial CF airway infection model following treatment with various tobramycin-glass combinations. Models were treated for 48 hours with bioactive glass extracts containing Ga^3+^ and/or Zn^2+^ ions at two dilutions (1/32, 1/16), with or without 16 µg/ml tobramycin. Bacterial growth was normalized to the untreated control for each bacteria strain. Data are presented as mean ± SEM (N = 2). Statistical analysis was performed using two-way ANOVA with Šidák’s multiple comparisons test; * p < 0.05, ** p < 0.01, *** p < 0.001, **** p < 0.0001.

When vancomycin-glass combinations were added to the polymicrobial CF airway infection model, *P. aeruginosa* growth was significantly decreased compared to the no treatment control in the presence of the Ga^3+^ or the Ga^3+^/Zn^2+^ glass (Figure 5). No significant change in *P. aeruginosa* growth occurred in the presence of the Zn^2+^ glass (Figure 5). *S. aureus* growth in the polymicrobial model was increased in the presence of the Ga^3+^/Zn^2+^ glass compared to the no treatment control, while the Ga^3+^ and the Zn^2+^ glasses did not produce a significant change in *S. aureus* growth (Figure 5).

**Figure 5.**
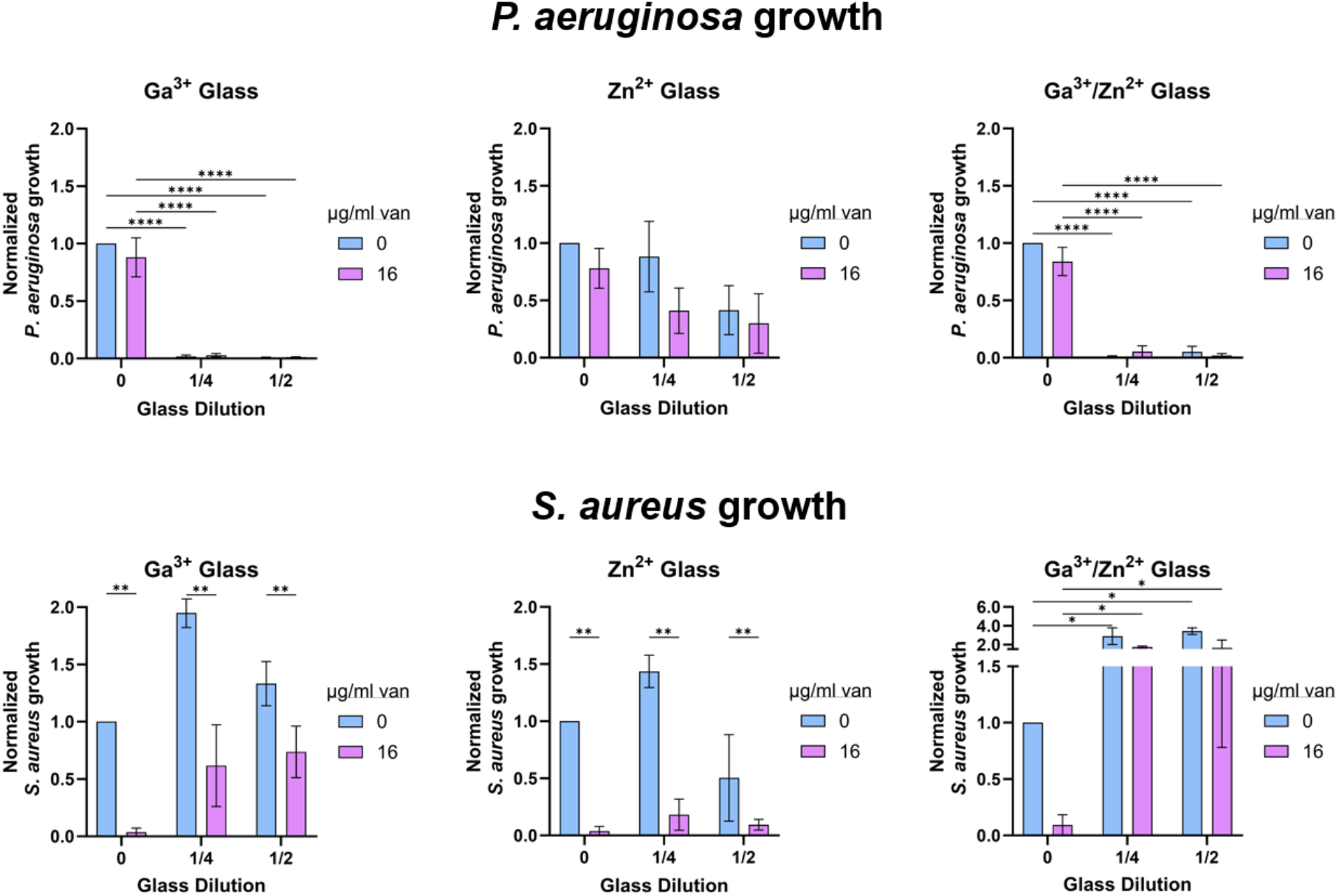
*P. aeruginosa* growth and *S. aureus* growth in the hydrogel phase of the polymicrobial CF airway infection model following treatment with various vancomycin-glass combinations. Models were treated for 48 hours with bioactive glass extracts containing Ga^3+^ and/or Zn^2+^ ions at two dilutions (1/4, 1/2), with or without 16 µg/ml vancomycin. Bacterial growth was normalized to the untreated control for each bacteria strain. Data are presented as mean ± SEM (N = 2). Statistical analysis was performed using two-way ANOVA with Šidák’s multiple comparisons test; * p < 0.05, ** p < 0.01, *** p < 0.001, **** p < 0.0001.

### Antibiotic-glass combinations do not impact mammalian cell viability in the airway infection model

Each antibiotic (tobramycin or vancomycin) was combined with a single bioactive glass extract (Ga^3+^, Zn^2+^, or Ga^3+^/Zn^2+^) and added to the CF airway infection model inoculated with either *P. aeruginosa* or *S. aureus* for a total of 48 hours. Following treatment, bronchial epithelial cell viability was determined using a live/dead assay.

In the *P. aeruginosa* model treated with tobramycin-glass combinations for 48 hours, all groups containing tobramycin showed high 16-HBE viability, exceeding 90% (Figure 6). In contrast, groups that received no tobramycin and no glass extracts had no visible cells remaining attached to the insert surface, and cell viability was therefore recorded as 0%—likely due to extensive *P. aeruginosa* overgrowth leading to widespread epithelial cell death. Groups treated with Ga^3+^ or Ga^3+^/Zn^2+^ glass, with or without tobramycin, also maintained 16-HBE viability above 90%, suggesting that these glasses did not negatively impact cell viability (Figure 6). However, in groups treated with Zn^2+^ glass alone, no epithelial cells remained, mirroring the outcome observed in the no treatment controls, indicating insufficient protection from bacterial overgrowth-induced cell death (Figure 6).

**Figure 6.**
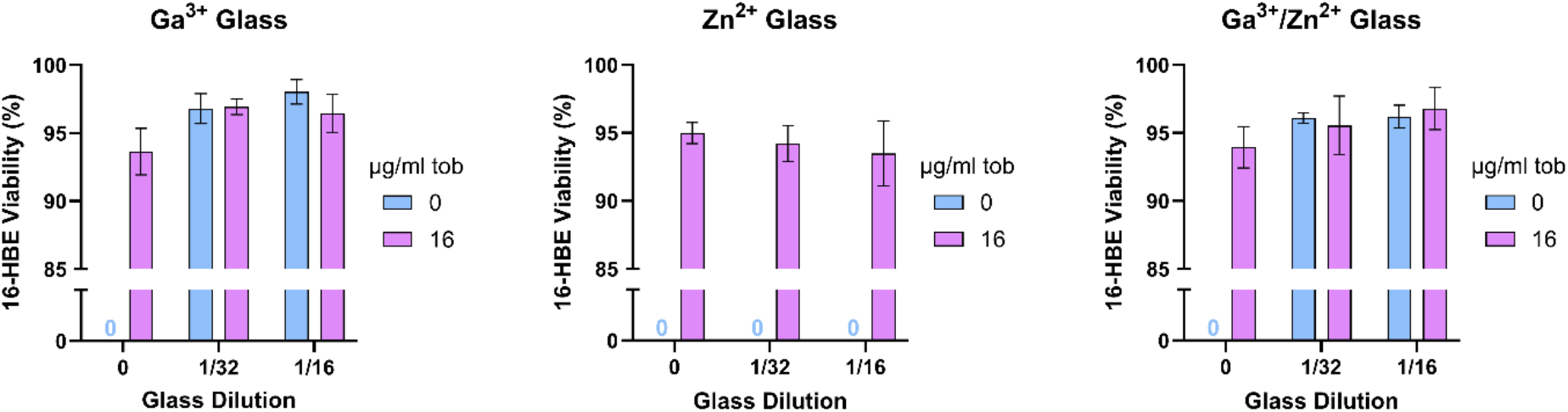
16-HBE viability in the CF airway infection model following treatment with various tobramycin-glass combinations. Models were treated for 48 hours with bioactive glass extracts containing Ga^3+^ and/or Zn^2+^ ions at two dilutions (1/32, 1/16), with or without 16 µg/ml tobramycin. Data are presented as mean ± SEM (N = 3).

In the *S. aureus* model treated with vancomycin-glass combinations for 48 hours, all groups lacking glass extracts—regardless of whether they received tobramycin or not—maintained 16-HBE viability above 90% (Figure 7). Similarly, treatment with any of the three glass extracts, with or without vancomycin, did not compromise cell viability, as 16-HBE viability consistently remained above 90% across all conditions (Figure 7).

**Figure 7.**
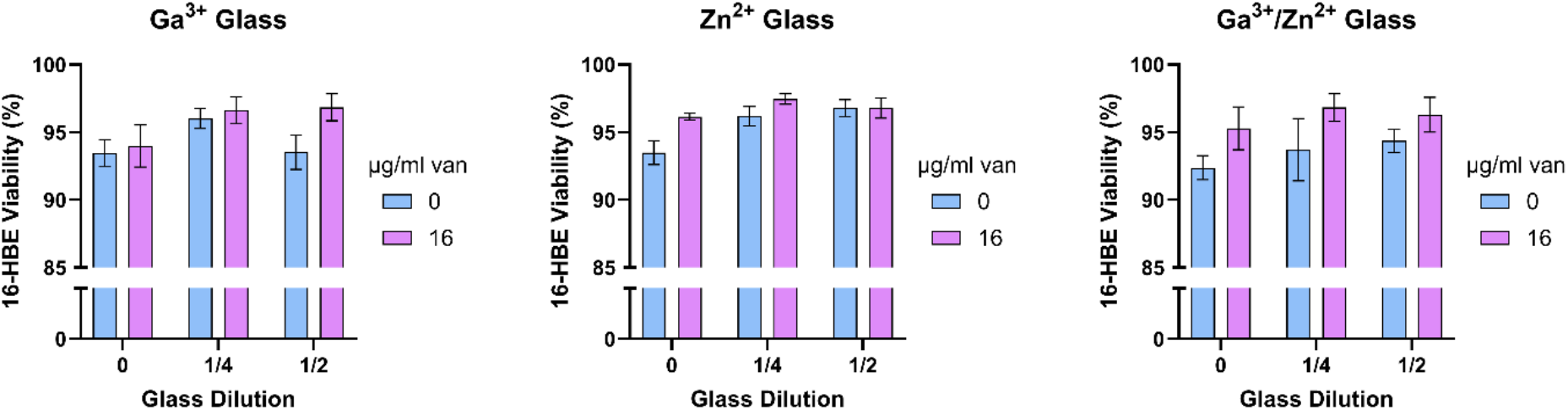
16-HBE viability in the CF airway infection model following treatment with various vancomycin-glass combinations. Models were treated for 48 hours with bioactive glass extracts containing Ga^3+^ and/or Zn^2+^ ions at two dilutions (1/4, 1/2), with or without 16 µg/ml vancomycin. Data are presented as mean ± SEM (N = 3).

## Discussion

Chronic microbial infections leading to lung damage and respiratory failure are the main causes of death in CF patients [1], [7]. The thick, dehydrated mucus in CF airways—caused by impaired ion transport from a genetic mutation—impairs mucociliary and cough clearance, creating a favorable environment for bacterial colonization [3], [5], [6]. The mucus also hinders antibiotic diffusion, making infections harder to treat and promoting antibiotic resistance [27]. These challenges highlight the need for novel therapies that can effectively suppress infection while limiting resistance. This study explores a novel combination therapy for CF airway infections using conventional antibiotics paired with metal ion-releasing bioactive glasses. These combinations were evaluated in an *in vitro* CF airway model to determine if bioactive glasses could enhance antibiotic efficacy through multiple antibacterial mechanisms.

The antibacterial efficacy of the bioactive glass extracts (Ga^3+^, Zn^2+^, Ga^3+^/Zn^2+^) against *P. aeruginosa* and *S. aureus* in a monomicrobial CF airway infection model were assessed over 48 hours. Both Ga^3+^-containing glasses significantly suppressed *P. aeruginosa* growth, whereas the Zn^2+^-only glass had no effect on *P. aeruginosa* growth (Figure 2). This outcome aligns with previous findings showing gallium is effective against bacteria reliant on iron metabolism, including *P. aeruginosa*, by disrupting iron-dependent pathways [19], [28]. Iron uptake in *P. aeruginosa* plays a key role in quorum sensing and virulence gene activation, and during chronic CF infections, *P. aeruginosa* will upregulate iron acquisition mechanisms to support growth [29], [30]. Due to its chemical similarity to the Fe^3+^ ion—including comparable octahedral and tetrahedral ionic radii—the Ga^3+^ ion acts as an iron mimic, binding iron-dependent sites and interfering with iron-dependent processes [31], [32]. This may explain the observed growth suppression as gallium competitively inhibits iron uptake, disrupting essential metabolic functions and leading to *P. aeruginosa* death. Additionally, gallium has demonstrated antibiofilm activity against Gram-negative bacteria like *P. aeruginosa* and *Acinetobacter baumannii* [33], [34]. In contrast, the Zn^2+^-only glass did not suppress *P. aeruginosa* growth in the CF airway infection model (Figure 2), which was unexpected given zinc’s known antibacterial properties across various species [35]. Zinc ions can disrupt bacterial metabolism by inhibiting glycolysis, increasing membrane permeability, generating ROS, and exerting antibiofilm effects [35], [36], [37]. A possible explanation is that the zinc concentration in the model was too low to trigger these effects, and higher concentrations might inhibit *P. aeruginosa* growth through one or more of these mechanisms. However, zinc is also essential for *P. aeruginosa* growth and virulence [38], and therefore at low concentrations, zinc may have instead supported bacterial survival by fulfilling its physiological roles, rather than acting as an antimicrobial agent.

As for *S. aureus* growth, both the Ga^3+^-only and Zn^2+^-only glasses significantly suppressed *S. aureus* growth, while the Ga^3+^/Zn^2+^ glass did not (Figure 3). The effect of the Ga^3+^-only glass was expected, as gallium disrupts iron metabolism and has some antibiofilm activity [31]. Like *P. aeruginosa, S. aureus* depends on iron for growth and colonization, so gallium may inhibit *S. aureus* by interfering with its iron-dependent pathways [39]. The Zn^2+^-only glass effectively suppressed *S. aureus* growth, consistent with zinc’s known antibacterial mechanisms—such as glycolysis inhibition, membrane disruption, and ROS generation [35], [40]. Zinc-based particles have also been shown to be particularly effective against Gram-positive bacteria like *S. aureus* [41]. The lack of antibacterial activity from the Ga^3+^/Zn^2+^ glass against *S. aureus* was unexpected, potentially indicating an antagonistic interaction between the two ions. One possibility is that zinc interferes with gallium uptake by competing with or blocking its substitution for iron, reducing gallium’s efficacy as an antibacterial agent. Alternatively, gallium and zinc may form poorly bioavailable metal complexes, lowering the concentration of active ions. This unexpected result highlights the need for further investigation.

Bacterial growth of *P. aeruginosa* and *S. aureus* in a monomicrobial CF airway infection model were assessed over 48 hours in the presence of bioactive glass extracts (Ga^3+^, Zn^2+^, Ga^3+^/Zn^2+^) combined with the antibiotics tobramycin and vancomycin, respectively. These experiments evaluated whether the glass extracts could enhance antibiotic efficacy. In the *P. aeruginosa* model, combining the Ga^3+^-only glass at the 1/32 and 1/16 dilutions with tobramycin significantly reduced *P. aeruginosa* growth compared to either treatment alone (Figure 2), indicating a synergistic effect. This likely results from the complementary mechanisms of action—gallium disrupts iron metabolism while tobramycin inhibits protein synthesis and damages the bacterial membrane [42], [43]. In contrast, the Ga^3+^/Zn^2+^ glass did not enhance tobramycin activity, suggesting no additive or synergistic effect. In the *S. aureus* model, the Zn^2+^-only glass at the 1/4 dilution combined with vancomycin significantly suppressed *S. aureus* growth more than either treatment alone (Figure 3). This may reflect the combined action of zinc, which generates ROS and disrupts metabolism, with vancomycin’s inhibition of cell wall synthesis [35], [44]. However, the Ga^3+^-only glass did not enhance vancomycin’s efficacy against *S. aureus*. Overall, these findings suggest that Ga^3+^-releasing glass can potentiate tobramycin against *P. aeruginosa*, while Zn^2+^-releasing glass can enhance vancomycin’s activity against *S. aureus* in a CF-like microenvironment.

*P. aeruginosa* and *S. aureus* were cultured together in the CF airway infection model and cultured with tobramycin-glass and vancomycin-glass combinations for 48 hours to assess antibacterial efficacy in a polymicrobial infection context. The Ga^3+^-containing glasses significantly reduced *P. aeruginosa* growth in the polymicrobial model (Figure 4; Figure 5), consistent with monomicrobial results, likely due to Ga^3+^ mimicking Fe^3+^ and disrupting iron metabolism [42]. In contrast, the Zn^2+^ glass had a minimal impact on *P. aeruginosa* growth (Figure 4; Figure 5), suggesting the zinc concentration was too low to induce antibacterial activity.

Notably, *P. aeruginosa* suppression by gallium was more pronounced in the polymicrobial model (Figure 4; Figure 5), likely due to interspecies competition between *P. aeruginosa* and *S. aureus*. As *P. aeruginosa* growth decreased, *S. aureus* likely expanded, further inhibiting *P. aeruginosa*. Supporting this, *S. aureus* growth increased up to 4-fold when treated with the Ga^3+^ glass, despite being suppressed by it in the monomicrobial model. This suggests that in the polymicrobial setting, relieving *P. aeruginosa*-mediated suppression outweighs gallium’s direct antibacterial effect on *S. aureus*. Similarly, the Zn^2+^ glass did not significantly reduce *S. aureus* growth in the polymicrobial model (Figure 4; Figure 5), differing from monomicrobial results. This may be due to dominant inhibition from *P. aeruginosa* masking any zinc-mediated effects on *S. aureus* growth.

Mammalian cell viability in the monomicrobial CF airway infection models was evaluated after 48 hours of exposure to the antibiotic-glass combinations. In the *P. aeruginosa* model, mammalian cell viability was preserved when 16 µg/ml tobramycin was added, but extensive cell death occurred in the absence of both tobramycin and glass (Figure 6). This suggests that *P. aeruginosa* overgrowth, when left untreated, leads to cytotoxicity. Supporting this, *P. aeruginosa* is known to induce host cell death via several mechanisms, including its Type III secretion system, virulence factors like pyocyanin (a ROS generator), and high motility [45], [46], [47]. In the tobramycin-glass treatment groups, mammalian cells remained viable with the Ga^3+^-containing glasses but not with the Zn^2+^-only glass (Figure 6). This aligns with the observation that Ga^3+^-containing glasses suppressed *P. aeruginosa* growth, while the Zn^2+^-only glass did not (Figure 2). Together, these results suggest that preventing *P. aeruginosa* overgrowth— particularly with gallium-based treatments—is critical to maintaining epithelial cell survival.

In contrast, in the *S. aureus* model, 16-HBE cells remained viable (>90%) even in the absence of vancomycin and glass (Figure 7), indicating that *S. aureus* overgrowth does not induce the same level of cytotoxicity. This may be due to differences in pathogenic mechanisms: *S. aureus* lacks ROS-generating metabolites and is non-motile [48], [49], which likely limits its spread and cytotoxic effects. Clinically, this mirrors findings where *P. aeruginosa* is more strongly associated with disease severity and mortality in CF compared to *S. aureus* [50].

This study presents a promising *in vitro* model for evaluating antibiotic-glass combinations in the context of CF airway infections; however, several limitations must be addressed to strengthen the model’s biological relevance and translational potential. One key limitation lies in the composition of the mucus-mimetic hydrogels used in the model. While these hydrogels successfully replicate the physical properties of healthy and CF mucus, they lack several biochemical components found in native airway secretions. Elements such as extracellular DNA, lipids, and non-mucin proteins contribute significantly to mucus structure and function *in vivo* [3]. Future work could explore the incorporation of these components into the hydrogel formulations to assess whether they influence bacterial behavior or alter the efficacy of antibiotic-glass treatments. Another limitation is the absence of a host immune component. The current model relies solely on airway epithelial cells and lacks the immune cells that play critical roles in infection progression and resolution in CF patients [51]. Incorporating immune cells such as macrophages or neutrophils would enable investigation of host-pathogen-therapy interactions and allow for a more comprehensive assessment of treatment efficacy in a CF-like microenvironment.

A further limitation relates to uncertainty surrounding the complete dissolution of the bioactive glasses used in this study. During preparation of the glass extract solutions, undissolved precipitates remained after 48 hours of incubation, suggesting that not all metal ions may have been released into solution. This could result in inconsistent concentrations of therapeutic ions such as Ga^3+^ and Zn^2+^ in the antibiotic-glass combinations. To ensure consistency and reproducibility, future work should include spectroscopic analysis to confirm the presence and concentration of metal ions in the extract solutions relative to their original glass formulations.

Looking ahead, future studies could focus on optimizing the glass formulations themselves. This could involve adjusting the type or concentration of antibacterial ions to more effectively target specific CF pathogens or altering the glass composition to better control the rate of ion release and improve solubility. The highly tunable nature of borate-based bioactive glasses makes them well-suited for such refinement [23]. Additionally, the efficacy of the developed antibiotic-glass combinations could be expanded to include other CF-associated pathogens, such as *Haemophilus influenzae, Stenotrophomonas maltophilia*, and *Aspergillus fumigatus* [1], [8]. The airway infection model also offers potential for adaptation to other mucosal infection contexts, such as the gastrointestinal or urogenital tracts.

## Conclusion

In this study, a novel therapeutic approach was developed to treat bacterial airway infections in CF by combining conventional antibiotics with metal ion-releasing bioactive glasses. Within a CF-like airway microenvironment, the growth of *P. aeruginosa* and *S. aureus* was effectively suppressed by the addition of these bioactive glass formulations, without compromising mammalian cell viability. Specific glass extracts demonstrated the ability to enhance the antibacterial activity of conventional antibiotics in monomicrobial CF airway infection models for both pathogens. In contrast, the polymicrobial co-infection model revealed more complex and distinct bacterial responses to the same antibiotic-glass combinations, highlighting the added challenge of treating polymicrobial infections. Notably, across all conditions where bacterial growth was significantly reduced, mammalian cell viability was preserved. These findings support the potential of antibiotic-bioactive glass combination therapies as a promising strategy for managing CF-related airway infections, with the added benefit of potentially mitigating the development of antibiotic resistance in CF-associated pathogens.

## Author Contributions

The manuscript was written through contributions of all authors. All authors have given approval to the final version of the manuscript. ‡These authors contributed equally. (match statement to author names with a symbol)

## Abbreviations

AA: antibiotic-antimycotic
ALG-MUC: alginate-mucin
ATPS: aqueoeus two-phase system
BHI: brain heart infusion
CaCl2: calcium chloride
CF: cystic fibrosis
CFTR: cystic fibrosis transmembrane conductance regulator
CFU: colony forming unit
DEX: dextran
DMEM: Dulbecco’s modified eagle medium
FBS: fetal bovine serum
LB: Luria-Bertani
PBS: phosphate-buffered saline
PEG: polyethylene glycol
ROS: reactive oxygen species

